# Attenuated Dopamine Receptor Signaling in Nucleus Accumbens Core in a Rat Model of Chemically-Induced Neuropathy

**DOI:** 10.1101/722801

**Authors:** D.E. Selley, M.F. Lazenka, L.J. Sim-Selley, D. N. Potter, Elena H. Chartoff, W.A. Carlezon, S.S. Negus

## Abstract

Neuropathy is major source of chronic pain that can be caused by mechanically or chemically induced nerve injury. Previous work in a rat model of neuropathic pain demonstrated that bilateral formalin injection into the hind paws produced mechanical hypersensitivity (allodynia) and depressed responding for intracranial self-stimulation (ICSS). To determine whether neuropathy alters dopamine receptor responsiveness in mesolimbic brain regions, we examined dopamine D_1_-like and D_2_-like receptor (D_1/2_R) signaling and expression in male rats 14 days after bilateral intraplantar formalin injections into both rear paws. D_2_R-mediated G-protein activation and expression of the D_2_R long, but not short, isoform were reduced in nucleus accumbens (NAc) core, but not in NAc shell, caudate-putamen (CPu) or ventral tegmental area (VTA) of formalin-compared to saline-treated rats. In addition, D_1_R-stimulated adenylyl cyclase (AC) activity was also reduced in NAc core, but not in NAc shell or prefrontal cortex, of formalin-treated rats, whereas D_1_R expression was unaffected. Expression of other proteins involved in dopamine neurotransmission, including dopamine uptake transporter (DAT) and tyrosine hydroxylase (TH), were unaffected by formalin treatment. In behavioral tests, the effects of D_2_R agonists on ICSS were attenuated in formalin-treated rats, whereas the effects of D_1_R agonists were unchanged. These results indicate that intraplantar formalin as a model of chemically induced neuropathy produces attenuation of highly specific DA receptor signaling processes in NAc core of male rats.

## 1. INTRODUCTION

The mesolimbic dopamine (DA) system consists of dopaminergic neurons that project from the ventral tegmental area (VTA) in midbrain to forebrain targets that include the nucleus accumbens (NAc) core and shell (Ikemoto et al., 2015; Sesack and Grace, 2010). Postsynaptic neurons in NAc consist primarily of two broad populations defined in part by their expression of either D_1_-like or D_2_-like DA receptors (D_1_R and D_2_R, respectively) (Beaulieu and Gainetdinov, 2011). Dopaminergic transmission between mesolimbic DA neurons and their postsynaptic NAc targets is known to play a major role in the neural representation of rewarding stimuli and the expression of motivated behavior, and a growing body of evidence suggests that pain states can impair that signaling (Mitsi and Zachariou, 2016; Taylor et al., 2016; Watanabe and Narita, 2018). As one example, we evaluated effects of intraperitoneal injection of dilute lactic acid (IP acid) in rats as an acute visceral noxious stimulus on both in vivo microdialysis measures of NAc core DA levels and intracranial self-stimulation (ICSS) as a positively reinforced operant behavior dependent on mesolimbic DA signaling (Leitl et al., 2014a). IP acid produced a concentration-dependent decrease in both NAc DA levels and ICSS, and both effects could be blocked by clinically effective analgesics, including morphine and ketoprofen. Manipulations often used in rodent models of chronic pain produce more subtle effects on mesolimbic DA signaling (Mitsi and Zachariou, 2016; Taylor et al., 2016; Watanabe and Narita, 2018) and behavior (Negus, 2019). However, we found previously that intraplantar (Ipl) injection of formalin served as a model of chemically induced neuropathy in rats that produced sustained changes in both a conventional metric of pain-related behavior (hypersensitive paw withdrawal from tactile stimuli delivered via von Frey filaments) as well as small but statistically significant and sustained depression of ICSS suggestive of sustained depression of mesolimbic DA signaling (Leitl and Negus, 2016; Leitl et al., 2014a). Microdialysis assessment of extracellular DA levels as a measure of DA signaling is best suited to within-subject evaluation of relatively transient changes in DA release over the course of minutes to hours, for example in studies with acute noxious or antinociceptive stimuli (Leitl et al., 2014a; Navratilova et al., 2012). More sustained changes in DA signaling, such as that potentially produced by chronic pain states, might be expected to alter more durable components of DA signaling, such as the activity or expression of DA receptors. Accordingly, the present study evaluated effects of Ipl formalin as a chronic neuropathic-pain stimulus on function and expression of DA receptors and related biomarkers of DA signaling.

Pain states have been associated with alterations in striatal dopamine D_2_R in both preclinical animal models and human imaging studies (Mitsi and Zachariou, 2016; Taylor et al., 2016). Human imaging studies have shown either increased or decreased striatal D_2_R receptor binding potentials using positron emission tomography in various pain states (Martikainen et al., 2018; Martikainen et al., 2015); however, these results could be due to changes in dopamine (DA) occupancy or altered D_2_R B_max_ levels (or both). In addition, the direction and magnitude of changes in D_2_R binding potentials can rapidly change in chronic pain patients in response to acute pain challenge (DaSilva et al., 2017; Martikainen et al., 2018). Moreover, human genetic studies have shown that polymorphism in the Drd2 gene is associated with baseline pain sensitivity, analgesic response to transcranial magnetic stimulation and likelihood of developing post-injury neuropathic pain (Jaaskelainen et al., 2014). Altogether, these findings suggest important roles for D_2_R function in the brain in human responses to painful stimuli.

Preclinical studies generally indicate decreases in D_2_R levels following chronic pain models in rodents, whereas synaptic DA levels have been reported to either increase or decrease. For example, dopaminergic neuronal burst firing in VTA and intra-NAc DA levels were increased, whereas D_2_R and tyrosine hydroxylase (TH) protein levels were decreased, two weeks after spared nerve injury (SNI) in rats (Sagheddu et al., 2015). Moreover, both D_2_R and D_1_R mRNA levels were decreased in NAc 28 days after surgery in this same SNI model (Chang et al., 2014). In contrast, NAc DA levels were decreased two weeks after peripheral nerve injury in mice (Taylor et al., 2014). Furthermore, decreased DA and increased dopamine transporter (DAT) levels were seen in NAc of rats two weeks after SNI (Wu et al., 2014). However, there are currently no published reports on DA receptor signaling at the biochemical level along with comprehensive measures of levels of DA receptors and other proteins associated with DA neurotransmission, such as DAT and TH. Here we examined the effects of Ipl formalin treatment of male rats on D_2_- and D_1_-like receptor-mediated G-protein signaling and protein levels of D_2_R, D_1_R and other proteins involved in regulating dopaminergic neurotransmission (DAT and TH) in mesolimbic dopaminergic brain regions, including NAc core and shell, prefrontal cortex (PFC) and VTA. Results of these experiments indicate attenuation in both D_2_R and D_1_R signaling in NAc core, so ICSS studies were then conducted to determine whether these changes were associated with altered ability of D_2_- and D_1_-like agonists to affect ICSS responding (Lazenka et al., 2016).

## 2. MATERIALS AND METHODS

### 2.1 Experimental Subjects

Adult male Sprague-Dawley rats (ENVIGO, Frederick, MD) were used for these studies. All rats had ad libitum access to food and water and were housed individually at Virginia Commonwealth University on a 12 hr light-dark cycle (6am – 6pm, lights on) in a facility accredited by the Association for the Assessment and Accreditation of Laboratory Animal Care. For molecular studies, rats weighed between 300 and 400 g at the time of Ipl injection. For ICSS, rats weighed between 300 and 400 g at the time of surgery to implant stimulating electrodes. All experiments were performed with the approval of the Institutional Animal Care and Use Committee at Virginia Commonwealth University in accordance with the *National Institutes of Health Guide for the Care and Use of Laboratory Animals 8^th^ edition* (National Research Council (U.S.), 2011).

### 2.2 In vitro functional assays: agonist-stimulated [^35^S]GTPγS binding and adenylyl cyclase activity

#### 2.2.1 Dissections

Fourteen days after Ipl treatment with formalin or saline, rats were euthanized by rapid decapitation, and brains were dissected using anatomical landmarks similar to those described previously in mice (Lazenka et al., 2017b; Wiebelhaus et al., 2015) with necessary modification. The prefrontal cortex was dissected by making a cut at the posterior extent of the anterior olfactory nucleus after which the olfactory nuclei were removed. This sample included frontal association, primary and secondary motor, anterior cingulate, prelimbic and orbital frontal cortices. A cut was then made anterior to the optic chiasm, producing a slice that included both the caudate-putamen and NAc. The nucleus accumbens was isolated by removing the cortex ventrally, the septum and nucleus of the horizontal limb of the diagonal band medially and separating the remaining tissue inferior to the caudate-putamen. The NAc core was separated from the NAc shell by removing tissue ventral, lateral and medial to the anterior commissure (shell) such that a tear-shaped structure tapering at the lateral ventricle superiorly and containing the anterior commissure was left (core). The remaining caudate-putamen was isolated by removing the corpus callosum and surrounding cortex. To dissect the VTA, slice was taken by first cutting immediately anterior to the mammillary bodies and a second cut was made anterior to the middle cerebellar peduncle, at the midpoint of the cerebral peduncles. From this slice, the interpeduncular nucleus/mammillary bodies, located ventrally; the substantia nigra, located laterally; and the interfascicular nucleus, located medially were removed. The remaining region ventral to the red nucleus was dissected and comprised primarily the VTA. All dissected tissue was stored at −80°C until use.

#### 2.2.2 Membrane preparation

Membranes were prepared from dissected brain regions as previously described (Lazenka et al., 2015). Briefly, tissue was thawed in membrane buffer (Tris-HCl, pH 7.4, 3 mM MgCl_2_, 1 mM EGTA), homogenized and centrifuged at 40,000 x g for 10 min. The pellet was collected and homogenized in assay buffer (Tris-HCl, pH 7.4, 3 mM MgCl_2_, 0.1 mM EGTA, 100 mM NaCl) and protein was determined by the Bradford method.

#### 2.2.3 [^35^S]GTPγS binding

Membranes were pretreated with adenosine deaminase (AD) for 15 min at 30°C prior to assay. [^35^S]GTPγS binding was conducted essentially as previously described (Lazenka et al., 2015). Briefly, membranes (3-6 µg protein were then incubated with varying concentrations of quinelorane (D_2_-like agonist) or CP55,940 (cannabinoid agonist), 30 µM GDP, and 0.1 nM [^35^S] GTPγS in assay buffer containing 0.1% BSA, for 2 hr at 30°C in a 0.5 ml total volume. Basal binding was assessed without agonist and nonspecific binding was measured with 20 µM unlabeled GTPγS. The assay was terminated by filtration through GF/B glass fiber filters, followed by 3 washes with ice-cold Tris buffer. Bound radioactivity was determined by liquid scintillation spectrophotometry.

#### 2.2.4 Adenylyl cyclase activity

Adenylyl cyclase assays were performed as previously described (Sim-Selley et al., 2011) with minor modifications. Membranes (20 µg protein) were pretreated with AD as above and incubated with varying concentrations of SKF82958 (D_1_-like agonist) or CGS21689 (A_2a_ agonist) in assay buffer containing 0.1% BSA, 50 µM ATP, 50 µM GTP, 0.2 mM DTT, 0.2 mM papaverine, 5 mM phosphocreatine, and 20 U/ml creatine phosphokinase for 15 min at 30°C. The incubation was terminated by addition of ELISA sample diluent, and cAMP was quantified using a cAMP ELISA kit (Arbor Assays, Ann Arbor, MI).

#### 2.2.5 Data analysis for in vitro functional assays

Data are reported as mean values ± SEM of 5-6 rats per group with samples assayed in duplicate. Concentration-effect curves were subjected to non-linear regression analysis to determine E_max_ and EC_50_ values. To control for day-to-day interassay variability, E_max_ values in both groups were also normalized to the saline-treated value on each day of the experiment. Significance of agonist-stimulation and formalin treatment was determined by two-way ANOVA with agonist concentration and formalin-versus-saline treatment as the main factors. Differences in E_max_ and EC_50_ values between groups were determined by the two-tailed Student’s *t*-test. All curve-fitting and statistical analysis was conducted using Prism software (GraphPad, San Diego, CA).

### 2.3 Protein quantification by immunoblot

#### 2.3.1 Dissections

Fourteen days after Ipl treatment with formalin or saline, rats were euthanized by rapid decapitation, and whole brains were frozen in isopentane at −30 °C and shipped frozen on dry ice to McLean Hospital for evaluation using methods similar to those described previously (Der-Avakian et al., 2017; Yap et al., 2015). Briefly, frozen brains were coronally sectioned on a cryostat (HM 505 E; Microm; Walldorf, Germany) until the following regions were exposed: NAc Core (Bregma 2.52mm), NAc Shell (Bregma 2.52mm), and VTA (Bregma −5.04mm), based on the atlas of Paxinos and Watson. Bilateral tissue punches 1–1.5 mm in length were taken with a 1 mm internal diameter corer (Fine Science Tools; Foster City, CA) and placed in Eppendorf tubes kept on dry ice and then stored at −80° C. After removal of the tissue cores, coronal sections (30 μM) of the exposed face of the brain were taken and Nissl stained with cresyl violet for histological analysis of placements. Only the rats in which the tissue punches were targeted appropriately were included in analyses.

#### 2.3.2 Immunoblotting

Tissue was sonicated in 1% sodium dodecyl sulfate (SDS) to dissociate membranes. Total protein concentrations in samples were determined using the Bio-Rad DC Protein Assay kit (Hercules, CA), and the concentration of each sample was adjusted to 2.0 mg/ml protein. Cell lysates were heated to 70°C for 10 min before polyacrylamide gel electrophoresis. Denatured protein (10-20 µg) was loaded per lane on 4-12% Bis-Tris Gels (Invitrogen). Protein was transferred onto PVDF membrane. Membranes were stained with Ponceau S then rapidly imaged for total protein as loading control/normalization before blocking for 2 hours at RT with 5% nonfat dry milk in TBS-T with 0.02% Tween 20, then incubated with mouse monoclonal anti-D_2_R (1:250; Santa Cruz SC-5303) in TBS-T overnight at 4C to detect long and short forms of D_2_R at 51 and 48kDa. Other primary antibodies included pTH (cell Signaling S3; 1; 1:1000); TH (Millipore/Sigma AB152; 1:40000); DAT (Millipore AB2231; 1:20000); D_1_R (Santa Cruz SC 14001; 1:250).

#### 2.3.2 Data analysis

Data are reported as mean values ± SEM of 5-6 rats per group. Immunoblots were analyzed by normalization of optical densities of immunoreactive protein to Ponceau stain, and then normalization of both groups to the saline condition. Significant differences were determined using Student’s *t*-test, conducted with Prism software.

### 2.4 In vivo assessment of intracranial self-stimulation (ICSS)

#### 2.4.1 Drugs for in vivo delivery

Formalin and (-)-quinpirole HCl (D_2_-like agonist) were obtained from Fisher Scientific (Waltham, MA) and Sigma-Aldrich (St. Louis, MO), respectively. (±)SKF82958 HBr [(±)-6-Chloro-7,8-dihydroxy-3-allyl-1-phenyl-2,3,4,5-tetrahydro-1H-3-benzazepine hydrobromide] was provided by the National Institute of Mental Health Chemical Synthesis and Drug Supply Program (Bethesda, MD). Each of the three compounds were dissolved in saline. Reagents used for molecular studies are described above. For all studies, rats received bilateral Ipl injections of 5% formalin (100 ul in each hind paw) or saline, as described previously (Leitl et al., 2014b). For behavioral studies, quinpirole and SKF82958 were administered by intraperitioneal (i.p.) injection, and drug doses are expressed in units of the salt form above.

#### 2.4.2 Surgery

Rats were anesthetized with isoflurane (3% in oxygen; Webster Veterinary, Phoenix, AZ, USA) until unresponsive to toe-pinch prior to implantation of stainless steel electrodes (Plastics One, Roanoke, VA, USA). The cathode, which was 0.25 mm in diameter and covered with polyamide insulation except at the flattened tip, was implanted utilizing a stereotactic method into the left medial forebrain bundle (MFB) at the level of the lateral hypothalamus using previously published coordinates (2.8 mm posterior to bregma, 1.7 mm lateral to the midsagittal suture, and 8.8 mm ventral to the skull) (Leitl et al., 2014b). Three screws were placed in the skull, and the anode (0.125 mm diameter, un-insulated) was wrapped around one of the screws to act as a ground. Dental acrylic was used to secure the electrode to the screws and skull. Ketoprofen (5 mg/kg) was administered as a postoperative analgesic immediately and 24 hrs following surgery. Animals were allowed to recover for at least one week before ICSS training.

#### 2.4.3 Apparatus

Operant conditioning chambers consisted of sound-attenuating boxes containing modular acrylic and metal test chambers (29.2 cm X 30.5 cm X 24.1 cm) (Med Associates, St. Albans, VT). Each chamber had a response lever (4.5 cm wide, 2.0 cm deep, 3.0 cm above the floor), a 2-watt house light, three stimulus lights (red, yellow and green) centered 7.6 cm above the lever, and an ICSS stimulator. Bipolar cables routed through a swivel-commutator (Model SL2C, Plastics One) connected the stimulator to the electrode. MED-PC IV computer software controlled all programming parameters and data collection (Med Associates).

#### 2.4.4 Training

The behavioral procedure was similar to that described previously for studies with formalin (Leitl and Negus, 2016; Leitl et al., 2014b) and with direct dopamine agonists (Lazenka et al., 2017a; Lazenka et al., 2016). A house light was illuminated during behavioral sessions, and lever-press responding under a fixed-ratio 1 (FR1) schedule produced delivery of a 0.5 s train of square-wave cathodal pulses (0.1 ms per pulse) via the intracranial electrode. During brain stimulation, the stimulus lights over the lever were illuminated, and responding had no scheduled consequences. During initial 60 min training sessions, stimulation intensity was set at 150 µA, and stimulation frequency was set at 158 Hz. Stimulation intensity was then individually manipulated in each rat to identify an intensity that maintained reinforcement rates >30 stimulations/min (range of 120 µA-215 µA for rats in this study). Once an appropriate intensity was identified, changes in frequency were introduced during sessions consisting of three consecutive 10 min components, each of which contained 10 consecutive 60 s trials. The stimulation frequency was 158 Hz for the first trial of each component, and frequency decreased in 0.05 log unit steps during the subsequent nine trials to a final frequency of 56 Hz. Each trial began with a 10 s time-out period, during which responding had no scheduled consequences, and five non-contingent stimulations at the designated frequency were delivered at 1 s intervals during the last 5 s of the time out. During the remaining 50 s of each trial, responding produced both intracranial stimulation at the designated frequency and illumination of the lever lights under an FR1 schedule as described above. ICSS performance was considered to be stable when frequency-rate curves were not statistically different over three consecutive days of training as indicated by lack of a significant effect of ‘day’ in a two-way analysis of variance (ANOVA) with day and frequency as the main effect variables (see Data Analysis below). All training was completed within six weeks of surgery.

#### 2.4.5 Testing

Once ICSS baselines were established, testing began using a 16-day protocol. On Day 2 of this protocol, rats were treated bilaterally with Ipl formalin (N=12) or saline (N=12). On Day 1 (the day before formalin/saline treatment), and on Day 16 (14 days after formalin/saline treatment), rats were tested with cumulative doses of either quinpirole (0.0032-0.1 mg/kg) or SKF82958 (0.01-0.32 mg/kg) (N=6 formalin-treated rats and N=6 saline-treated rats for each drug). Cumulative-dosing studies with quinpirole or SKF82958 were accomplished using test sessions that consisted of three baseline components followed by four consecutive 30-min test periods. Each test period consisted of a 10 min time out followed by a pair of 10-min test components. A dose of quinpirole or SKF82958 was administered i.p. in a volume of 1 ml/kg at the start of each time out, and each dose increased the total cumulative dose by a half-log increment. Three-component ICSS baseline sessions were conducted on most other days (excluding weekends).

#### 2.4.6 Data analysis for ICSS

The first baseline component for each day was considered to be a “warm-up” component, and data were discarded. The primary dependent variable was reinforcement rate in stimulations per min during each frequency trial for all remaining baseline and test components. To normalize these data, raw reinforcement rates from each trial in each rat were converted to percent maximum control rate (%MCR) for that rat. The MCR was defined as the mean of the maximal rates observed during the second and third baseline components for the three consecutive training days preceding the 16-day test protocol (six total baseline components). Subsequently, % MCR values for each trial were calculated as [(reinforcement rate during a frequency trial)/(MCR)]×100. For each rat during each session, data from baseline and test components were averaged to yield baseline and test frequency-rate curves. Baseline and test data were then averaged across rats to yield mean baseline and test frequency-rate curves for each manipulation. Results were compared by repeated measures two-way ANOVA with ICSS frequency as a within-subject factor and either Ipl treatment (between-subjects factor) or dopamine agonist dose (within-subjects factor) as the second factor. A significant ANOVA was followed by the Holm-Sidak post-hoc test, and the criterion for significance was *p* < .05.

To provide an additional summary measure of ICSS performance, the total number of stimulations per component was determined across all 10 frequency trials of each component. To normalize these data, raw numbers of stimulations per component in each rat were converted to the percent baseline number of stimulations per component for that rat, with the baseline defined as the mean number of stimulations per component during the second and third components on the three consecutive baseline days preceding testing. Thus, % Baseline Stimulations was calculated as (mean total stimulations during a component/mean total stimulations during baseline components) x 100. These data were then averaged across rats. Baseline data in formalin- and saline-treatment groups on a given day were compared by t-test. Drug effects in formalin- and saline-treatment groups on a given day were compared by two-way mixed ANOVA with dose as a within-subjects factor and Ipl treatment as a between-subjects factor. A significant ANOVA was followed by the Bonferroni Holm-Sidak post-hoc test, and the criterion for significance was *p* < 0.05. All statistical analyses were conducted with Prism software.

## 3. RESULTS

### 3.1 Formalin treatment selectively attenuates D_2_-like receptor-mediated G-protein activation in NAc core

The effect of Ipl formalin treatment on G-protein activation by dopamine D_2_-like receptors was determined with quinelorane-stimulated [^35^S]GTPγS binding in membranes prepared from mesolimbic dopaminergic brain regions, including NAc core and shell, caudate-putamen (CPu) and VTA. Basal [^35^S]GTPγS binding varied among regions, but was unaffected by formalin treatment in any region examined, as indicated by two-way ANOVA (Supplementary Figure 1A). Concentration-effect curves of quinelorane-stimulated [^35^S]GTPγS binding were examined in NAc core versus shell. Results in NAc core showed decreased D_2_R-stimulated activity in formalin-relative to saline-treated-rats, as confirmed by two-way ANOVA, which revealed a main effect of formalin treatment (Figure 1A). Curve-fitting analysis revealed trends toward a significant decrease in E_max_ (t(8) = 1.995, p = 0.081) and increase in log EC_50_ [t(8) = 2.004, p = 0.080] values (Table 1). Normalization of E_max_ values in formalin-treated rats to each corresponding value in saline-treated rats revealed a significant decrease [t(8) = 2.514, p = 0.036] in formalin-compared to saline-treated rats (Table 1).

**Table 1.** E_max_ and EC_50_ values of agonist-stimulated [^35^S]GTPγS binding in formalin- and vehicle-treated rats

| Treatment | Agonist | E<sub>max</sub> (% Stim.) | E<sub>max</sub> (% Saline) | EC<sub>50</sub> (log M) |
| --- | --- | --- | --- | --- |
| <u>NAc Core:</u> |  |  |  |  |
| Saline | Quinelorane | 50.3 ± 6.1 | 100.0 ± 12.1 | -6.91 ± 0.04 |
| Formalin | Quinelorane | 32.0 ± 5.5 | 61.9 ± 6.2* | -6.72 ± 0.07 |
| Saline | CP55,940 | 103.1 ± 7.7 | 100.0 ± 7.5 | -8.22 ± 0.05 |
| Formalin | CP55,940 | 107.6 ± 7.0 | 105.0 ± 3.1 | -8.08 ± 0.06 |
| <u>NAcShell:</u> |  |  |  |  |
| Saline | Quinelorane | 31.1 ± 2.8 | 100.0 ± 9.1 | -6.95 ± 0.11 |
| Formalin | Quinelorane | 34.4 ± 2.9 | 114.0 ± 10.0 | -7.00 ± 0.08 |
| Saline | CP55,940 | 113.4 ± 4.3 | 100.0 ± 3.8 | -7.93 ± 0.06 |
| Formalin | CP55,940 | 116.8 ± 3.1 | 104.4 ± 4.0 | -7.99 ± 0.05 |
| <u>VTA</u> |  |  |  |  |
| Saline | Quinelorane | 10.2 ± 0.9 | 100.0 ± 9.1 | -7.03 ± 0.19 |
| Formalin | Quinelorane | 8.4 ± 1.3 | 88.4 ± 12.3 | -7.35 ± 0.16 |
| <u>CPu</u> |  |  |  |  |
| Saline | Quinelorane | 60.7 ± 4.2 | 100.0 ± 7.0 | -7.00 ± 0.17 |
| Formalin | Quinelorane | 67.3 ± 6.6 | 110.8 ± 10.8 | -6.95 ± 0.08 |

Concentration-effect curves of quinelorane- or CP55,940-stimulated [<sup>35</sup>S]GTPγS binding in NAc core, NAc shell, VTA and CPu were fit by non-linear regression analysis to obtain E<sub>max</sub> and EC<sub>50</sub> values. Values are presented as mean ± SEM (n = 5-6). % Stim: % stimulation; % Saline: E<sub>max</sub> values normalized to the maximal stimulation obtained in saline-treated mice on each day of assay.

**Figure 1.**
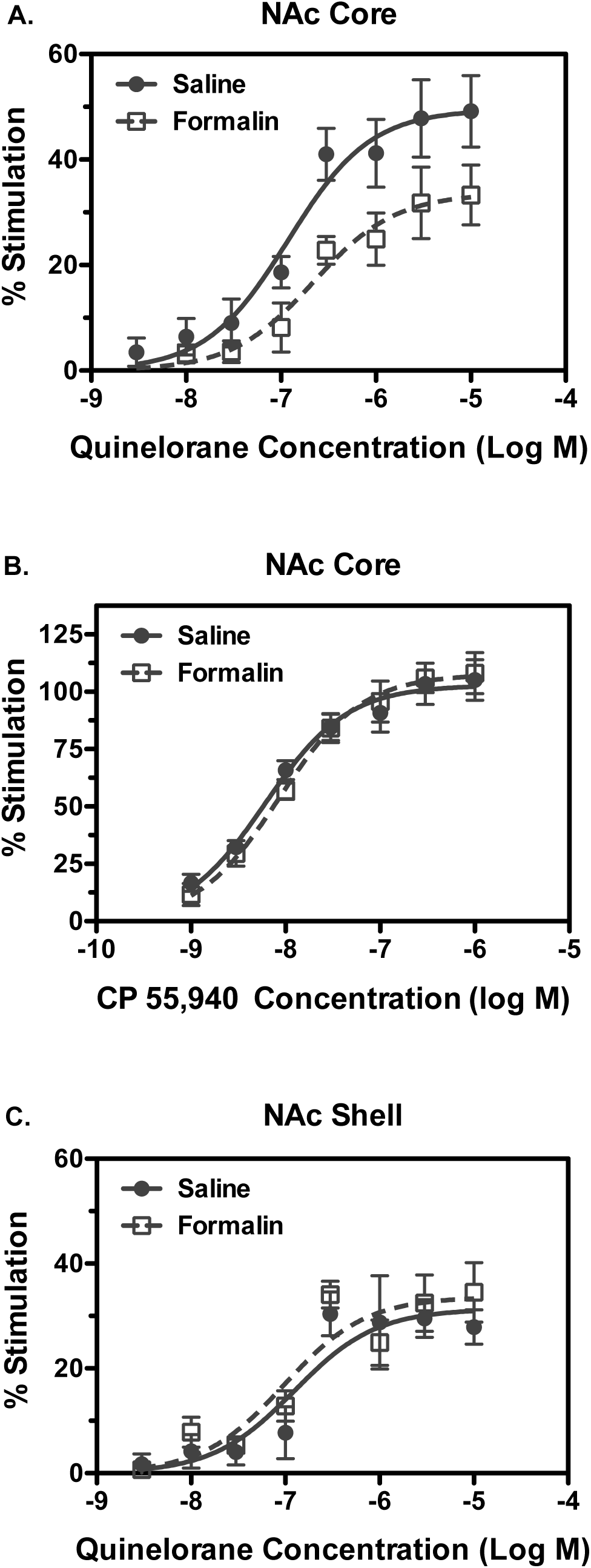
Formalin treatment decreases D_2_-like agonist-stimulated G-protein activity in NAc core but not shell. Concentration-effect curves were conducted for stimulation of [^35^S]GTPγS binding by quinelorane (A, C) or CP55,940 (B) in membranes from NAc core (A, B) or shell (C). Data are mean ± SEM of % stimulation of [^35^S]GTPγS binding (n = 5). Quinelorane-stimulated [^35^S]GTPγS binding was reduced in NAc core (A) but not shell (C) of formalin-relative to saline-treated rats. Two-way ANOVA in NAc core showed a main effect of qunelorane concentration [F(7, 59) = 23.05, p < 0.0001] and formalin treatment [F(1,59) = 20.73, p < 0.0001], whereas in NAc shell there was a main effect of qunelorane concentration [F(7,61) = 19.62, p < 0.0001] but not of formalin treatment, with no significant interactions between factors in either region. There was no difference between experimental groups in CP55,940-stimulated [^35^S]GTPγS binding in NAc core (B). Two-way ANOVA revealed a main effect of CP55,940 concentration [F(6,56) = 61.51, p < 0.0001] but not of formalin treatment nor was there an interaction.

To determine whether the formalin-induced decrease in G-protein activation was homologous to D_2_-like receptors, activity of the CB_1_ cannabinoid receptor, another G_i/o_-coupled receptor that has overlapping localization with D_2_ R in NAc (Pickel et al., 2006), was examined. In contrast to results with the D_2_-like agonist quinelorane, G-protein activation by the cannabinoid agonist CP55,940 was unaffected by formalin treatment, as indicated by two-way ANOVA (Figure 1B). E_max_ and log EC_50_ values of CP55,940 did not differ between formalin- and saline-treated rats (Table 1).

Formalin treatment did not alter D_2_R-mediated G-protein activation in NAc shell, as indicated by two-way ANOVA (Figure 1C). Quinelorane E_max_ and log EC_50_ values did not differ between formalin- and saline-treated rats (Table 1). Similarly, CP55,940-stimulated G-protein activation in NAc shell was unaffected by formalin treatment (data not shown). E_max_ and log EC_50_ values of CP55,940 did not differ between formalin- and saline-treated rats (Table 1).

D_2_-like receptor-stimulated G-protein activation was also assessed in VTA and CPu (Supplemental Figure 2). Results in VTA showed low and variable stimulation of [^35^S]GTPγS binding by quinelorane, which did not differ between saline- and formalin-treated rats, as indicated by two-way ANOVA. Accordingly, quinelorane E_max_ and log EC_50_ values in VTA did not differ between groups (Table 1). Quinelorane produced more robust stimulation in CPu, but there was no effect of formalin treatment, as indicated by two-way ANOVA. Quinelorane E_max_ and log EC_50_ values in CPu did not differ between saline- and formalin-treated rats (Table 1). Altogether, these results indicate that Ipl formalin treatment attenuated D_2_R-mediated G-protein activation in NAc core, but not in NAc shell, CPu or VTA, without affecting CB_1_ cannabinoid receptor-mediated G-protein activation.

### 3.2 Formalin treatment selectively attenuates D_1_-like receptor-mediated AC activation in NAc core

The effect of Ipl formalin treatment on G-protein signaling by dopamine D_1_-like receptors was determined with SKF82958-stimulated AC activity in membranes prepared from NAc core and shell. Basal AC activity varied among regions but was unaffected by formalin treatment, as indicated by two-way ANOVA (Supplemental Figure 1B). Concentration-effect curves of SKF82958-stimulated AC activity were examined in NAc core versus shell. Results in NAc core showed decreased D_1_R-stimulated activity in formalin-relative to saline-treated rats, as confirmed by two-way ANOVA, which showed a main effect of formalin treatment (Figure 2A). Curve-fitting analysis revealed a significant decrease in both E_max_ [t(12) = 2.308, p = 0.039] and normalized E_max_ values [t(12) = 2.869, p = 0.014] of SKF82958 in formalin-compared to saline-treated rats, with no difference in log EC_50_ values (Table 2). In contrast, AC activation in NAc core by the adenosine A_2_a agonist CGS21680 was unaffected by formalin treatment, as indicated by two-way ANOVA (Figure 2B). E_max_ and log EC_50_ values of CGS21680 did not differ between formalin- and saline-treated rats (Table 2).

**Figure 2.**
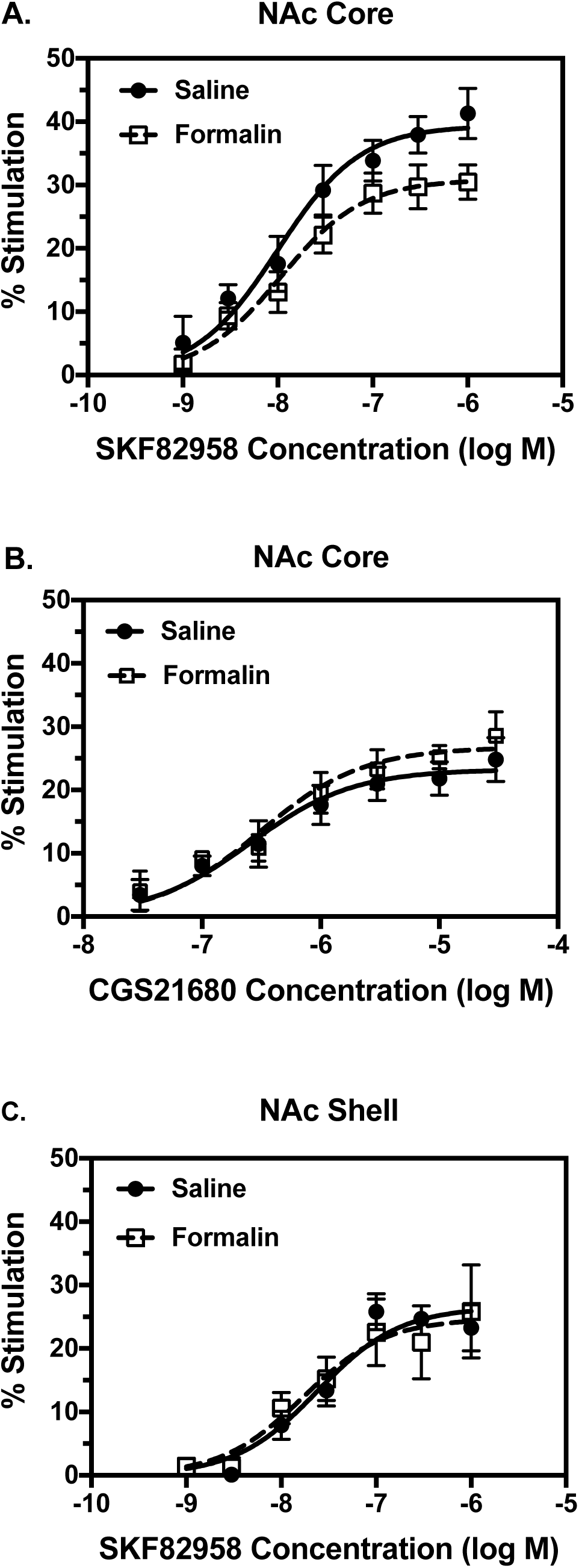
Formalin treatment selectively decreases D_1_-like agonist-stimulated AC activity in NAc core but not shell. Concentration-effect curves were conducted for stimulation of AC activity by SKF82958 (A, C) or CGS21680 (B) in membranes from NAc core (A, B) or shell (C). Data are mean ± SEM of % stimulation of AC activity (n = 5-7). SKF82958-stimulated AC activity was reduced in NAc core (A) but not shell (C) of formalin-relative to saline-treated rats. Two-way ANOVA in NAc core showed a main effect of quinelorane concentration [F(6, 75) = 29.45, p < 0.0001) and formalin treatment [F(1,75) = 11.63, p < 0.0001), where as in NAc shell there was a main effect of SKF82958 concentration [F(6,52) = 18.47, p < 0.0001] but not of formalin treatment, with no significant interactions between factors in either region. There was no difference between experimental groups in CGS21680-stimulated AC activity in NAc core (B). Two-way ANOVA revealed a main effect of CGS21680 concentration [F(6,70) = 17.25, p < 0.0001] but not of formalin treatment nor was there an interaction.

**Table 2.**
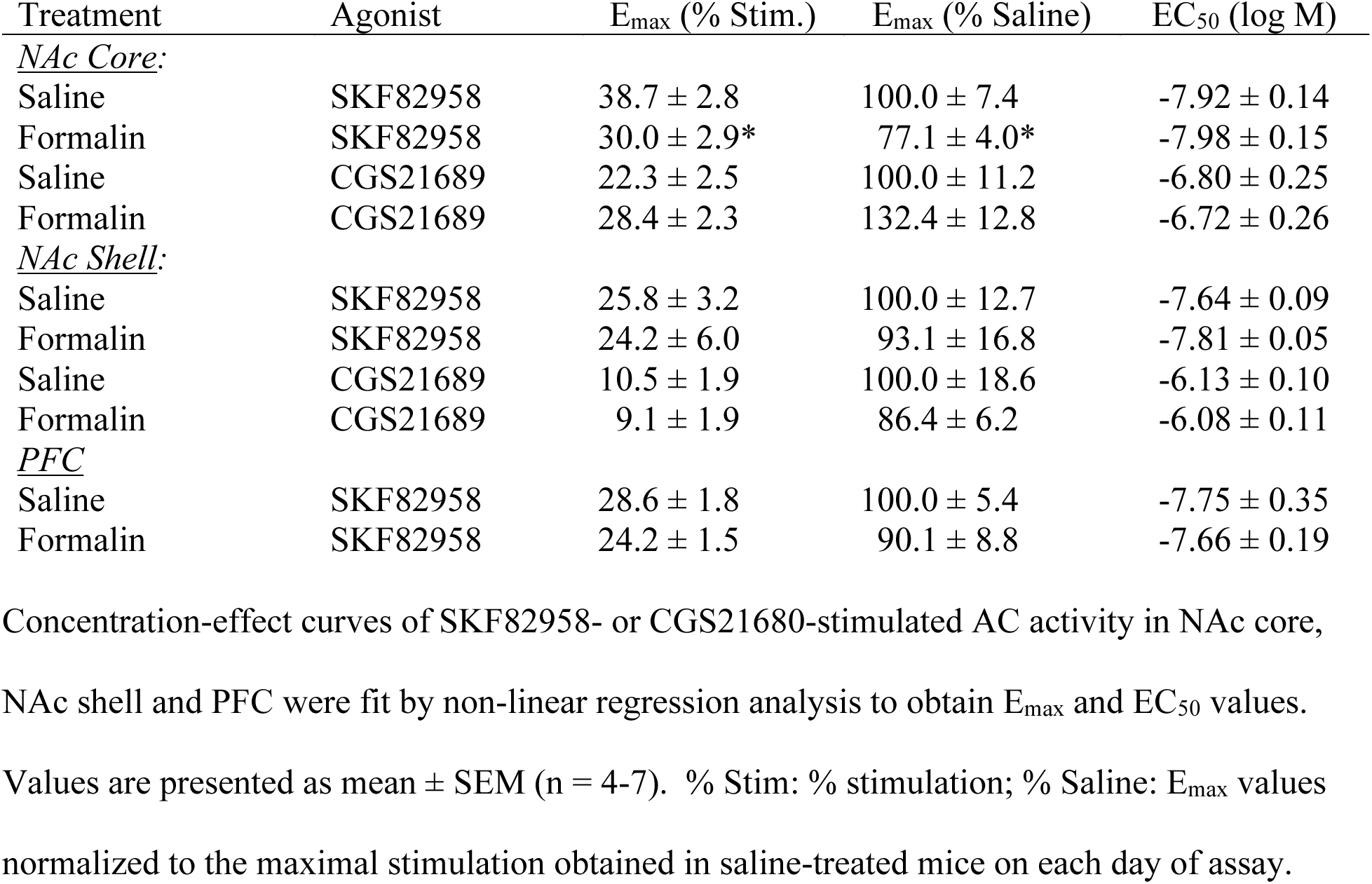
E_max_ and EC_50_ values of agonist-stimulated AC activity in formalin- and vehicle-treated rats

| Treatment | Agonist | E<sub>max</sub> (% Stim.) | E<sub>max</sub> (% Saline) | EC<sub>50</sub> (log M) |
| --- | --- | --- | --- | --- |
| <u>NAc Core:</u> |  |  |  |  |
| Saline | SKF82958 | 38.7 ± 2.8 | 100.0 ± 7.4 | -7.92 ± 0.14 |
| Formalin | SKF82958 | 30.0 ± 2.9* | 77.1 ± 4.0* | -7.98 ± 0.15 |
| Saline | CGS21689 | 22.3 ± 2.5 | 100.0 ± 11.2 | -6.80 ± 0.25 |
| Formalin | CGS21689 | 28.4 ± 2.3 | 132.4 ± 12.8 | -6.72 ± 0.26 |
| <u>NAc Shell:</u> |  |  |  |  |
| Saline | SKF82958 | 25.8 ± 3.2 | 100.0 ± 12.7 | -7.64 ± 0.09 |
| Formalin | SKF82958 | 24.2 ± 6.0 | 93.1 ± 16.8 | -7.81 ± 0.05 |
| Saline | CGS21689 | 10.5 ± 1.9 | 100.0 ± 18.6 | -6.13 ± 0.10 |
| Formalin | CGS21689 | 9.1 ± 1.9 | 86.4 ± 6.2 | -6.08 ± 0.11 |
| <u>PFC</u> |  |  |  |  |
| Saline | SKF82958 | 28.6 ± 1.8 | 100.0 ± 5.4 | -7.75 ± 0.35 |
| Formalin | SKF82958 | 24.2 ± 1.5 | 90.1 ± 8.8 | -7.66 ± 0.19 |

Concentration-effect curves of SKF82958- or CGS21680-stimulated AC activity in NAc core, NAc shell and PFC were fit by non-linear regression analysis to obtain E<sub>max</sub> and EC<sub>50</sub> values. Values are presented as mean ± SEM (n = 4-7). % Stim: % stimulation; % Saline: E<sub>max</sub> values normalized to the maximal stimulation obtained in saline-treated mice on each day of assay.

Formalin treatment did not alter D_1_R-mediated AC activation in NAc shell, as indicated by two-way ANOVA (Figure 2C). SKF82958 E_max_ and log EC_50_ values did not differ between formalin- and saline-treated rats (Table 2). CGS21680-stimulated AC activity was minimal (E_max_ ≤ 10%) in NAc shell and showed concentration-dependent stimulation only in 4 of the 5 sample pairs, so 4 samples per group were included in the analysis (data not shown). E_max_ and log EC_50_ values of CGS21680 did not differ between formalin- and saline-treated rats (Table 2).

Because D_1_ and A_2a_ receptors are the major G_s/olf_-coupled receptors driving AC activation in each of the two distinct populations of striatal medium spiny neurons, the effect of formalin on the ratio of maximal AC activation by each receptor was determined. The ratio of D_1_/A_2a_-stimulated AC activity was decreased by ∼41% in NAc core of formalin-relative to saline-treated rats (1.11 ± 0.16 versus 1.87 ± 0.28, respectively; [t(12) = 2.430, p = 0.032]). In contrast, formalin treatment did not affect the ratio of D_1_/A_2a_-stimulated AC activity in NAc shell (2.79 ± 0.46 versus 2.49 ± 0.20 in formalin- and saline-treated rats, respectively).

Formalin treatment also did not alter D_1_R-mediated AC activation in the prefrontal cortex (PFC), a dopaminergic terminal field region that is enriched in D_1_-like receptors (Herve et al., 1992) (Supplemental Figure 3). SKF82958 stimulated AC activity in a concentration-dependent manner, but there was no effect of formalin treatment, as indicated by two-way ANOVA. Neither E_max_ nor log EC_50_ values of SKF82958 differed between saline- and formalin-treated rats (Table 2). Taken together, these results indicate that formalin treatment attenuated D_1_R-mediated AC activation in NAc core, but not in NAc shell or PFC, without significantly altering adenosine A_2a_ receptor-mediated AC activation. Moreover, the ratio of D_1_:A_2a_-stimulated AC activity was significantly decreased in NAc core but not shell.

### 3.3 Formalin treatment selectively decreases D_2L_ receptor protein levels in NAc core

The receptor function experiments described above showed decreases in both D_2_-like receptor-mediated G-protein activation and D_1_-like receptor-mediated AC activation in NAc core but not shell of formalin-treated rats. Because decreased functional activity could be due to decreased receptor levels, reduced coupling between each receptor and its cognate G-protein or both, D_2_R and D_1_R protein levels were determined by immunoblot analysis of NAc core and shell. Results in NAc core (Figure 3A, C and E; Supplemental Figures 3 and 4) showed a ∼17% decrease in D_2_R long isoform (D_2L_) immunoreactivity in formalin-compared to saline-treated rats [t(10) = 2.363, p = 0.020], without any difference in D_2_R short isoform (D_2S_) levels.

**Figure 3.**
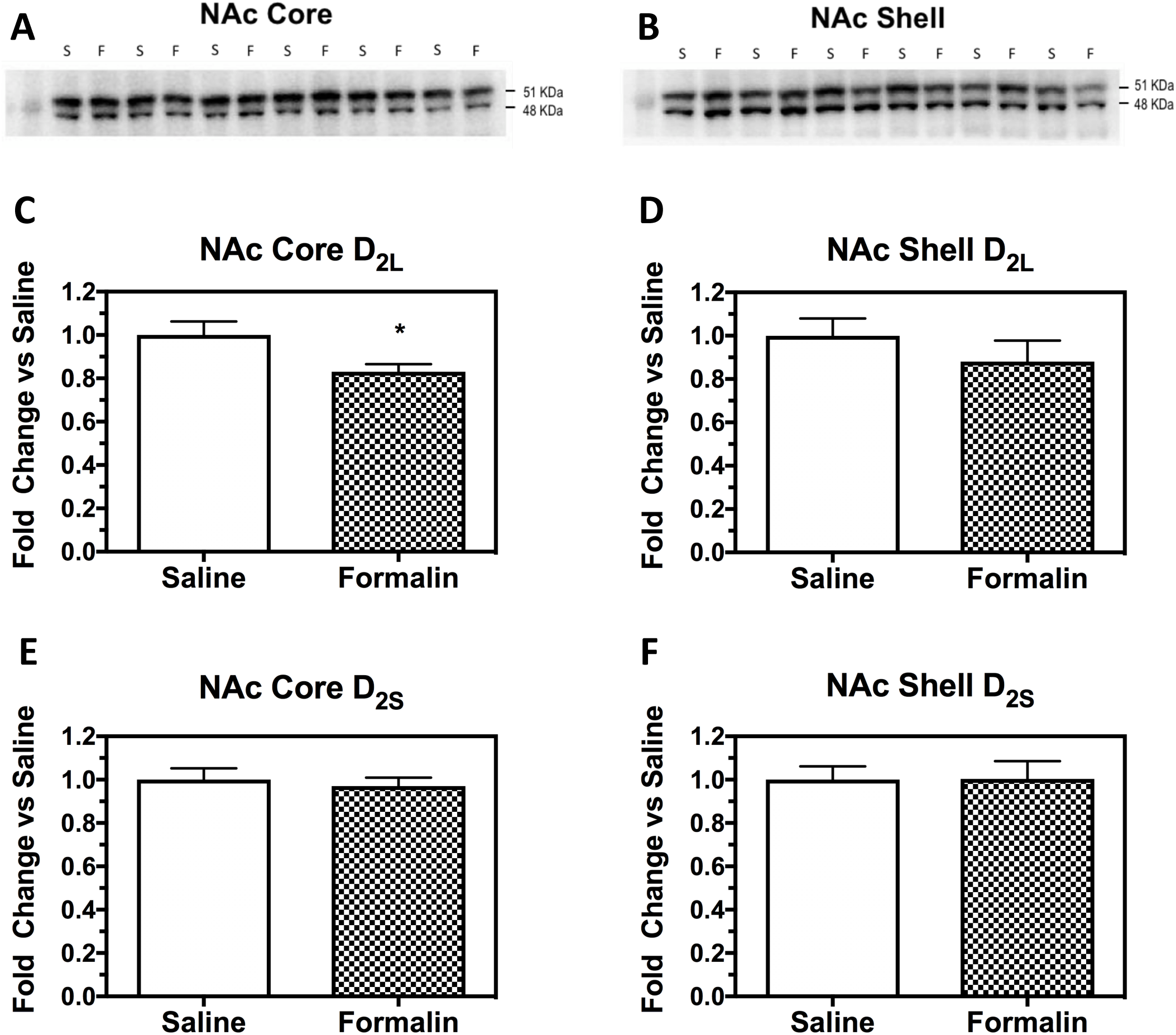
Formalin treatment decreases D_2L_ protein levels in NAc core but not shell. Immunoblots were conducted for D_2_ receptor protein in punches of NAc core (A, C, E) or shell (B, D, F). (A, B) Immunobot images of D_2_ receptor immunoreactivity. (C-F) Data are mean ± SEM of fold change in D_2L_ isoform (C, D) or D_2S_ isoform (E, F) in each sample relative to the mean value obtained in saline-treated rats (n = 6). Only the D_2L_ isoform in NAc core was significantly decreased in formalin-relative to saline-treated rats. * p < 0.05 by Student’s t-test.

In contrast to results in NAc core, no differences in either D_2L_ or D_2S_ receptor levels were seen in NAc shell (Figure 3B, D and F). D_1_R immunoreactivity also did not differ between saline- and formalin-treated rats in either NAc core or shell (Supplemental Figures 6 and 7). Likewise, D_2L_, D_2S_ and D_1_ receptor immunoreactivity did not differ between formalin- and saline-treated rats in VTA or PFC (data not shown). Furthermore, immunoblot analysis of other proteins important in the regulation of dopamine neurotransmission, including DAT tyrosine hydroxylase (TH) and phospho-TH, also showed no differences between formalin- and saline-treated rats in NAc core or shell, VTA or PFC (data not shown). Altogether, these results indicate that formalin treatment selectively decreased D_2L_R protein in the NAc core without affecting other proteins directly involved in dopamine neurotransmission.

#### 3.4 Formalin treatment attenuates modulation of ICSS responding by D_2_ but not D_1_ receptor activation

Under baseline conditions, MFB stimulation maintained a frequency-dependent increase in reinforcement rates. The mean ± SEM maximum control rate for all rats in the study was 53 ± 1.9 stimulations per trial, and the mean ± SEM baseline number of stimulations per component was 224 ± 11.5. There were no differences across groups in either baseline ICSS or in drug effects on ICSS on Day 1, before formalin or saline treatment (data not shown). Figure 4 shows that there was also no difference in baseline ICSS performance on Day 16, 14 days after formalin or saline treatment. Figure 5 shows effects of quinpirole and SKF82958 on Day 16 in each group. Quinpirole produced dose-dependent rightward/downward shifts in the ICSS frequency-rate curves in both groups; however, quinpirole was less potent to decrease the number of stimulations per component in the formalin-treated group than in the saline-treated group. SKF82958 produced a mixed profile of ICSS facilitation and ICSS depression in both groups. In particular, a dose of 0.1 mg/kg SKF82958 facilitated ICSS for at least one brain-stimulation frequency in both groups, and a higher dose of 0.32 mg/kg SKF82958 decreased ICSS for at least two brain-stimulation frequencies in both groups. There were no significant differences between formalin- and saline-treated groups in SKF82958 effects on the number of stimulations per component.

**Figure 4.**
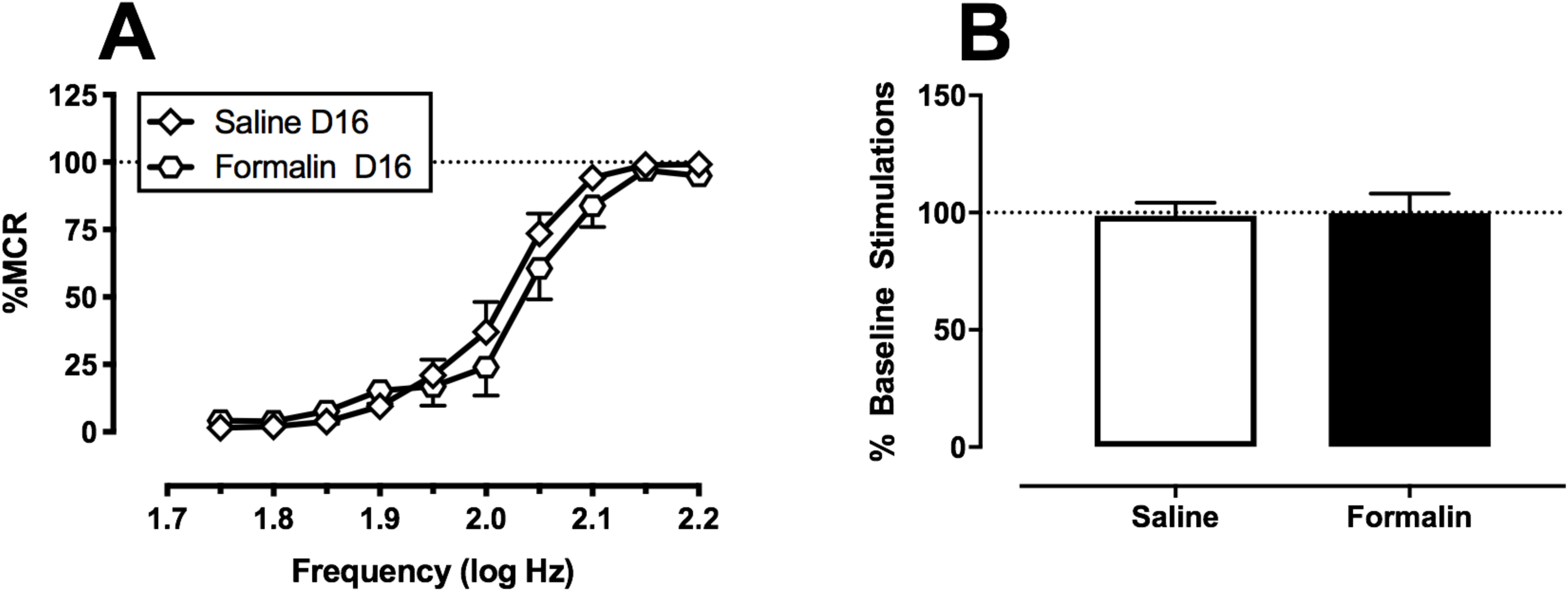
Baseline ICSS performance was unaffected by Ipl formalin treatment. Panel A: Abscissa: Frequency of electrical brain stimulation in Hz (log scale). Ordinate: Percent maximum control reinforcement rate (% MCR). Two-way ANOVA indicated a significant main effect of frequency [F(9, 198) = 151, p < 0.0001], but no effect of treatment or frequency x treatment interaction. Panel B: Abscissa: Intraplantar treatment group. Ordinate: Percent baseline number of stimulations per component, a summary measure of ICSS performance across all brain stimulation frequencies. T-test indicated no difference between treatment groups. All data show mean ± SEM of 12 rats.

**Figure 5.**
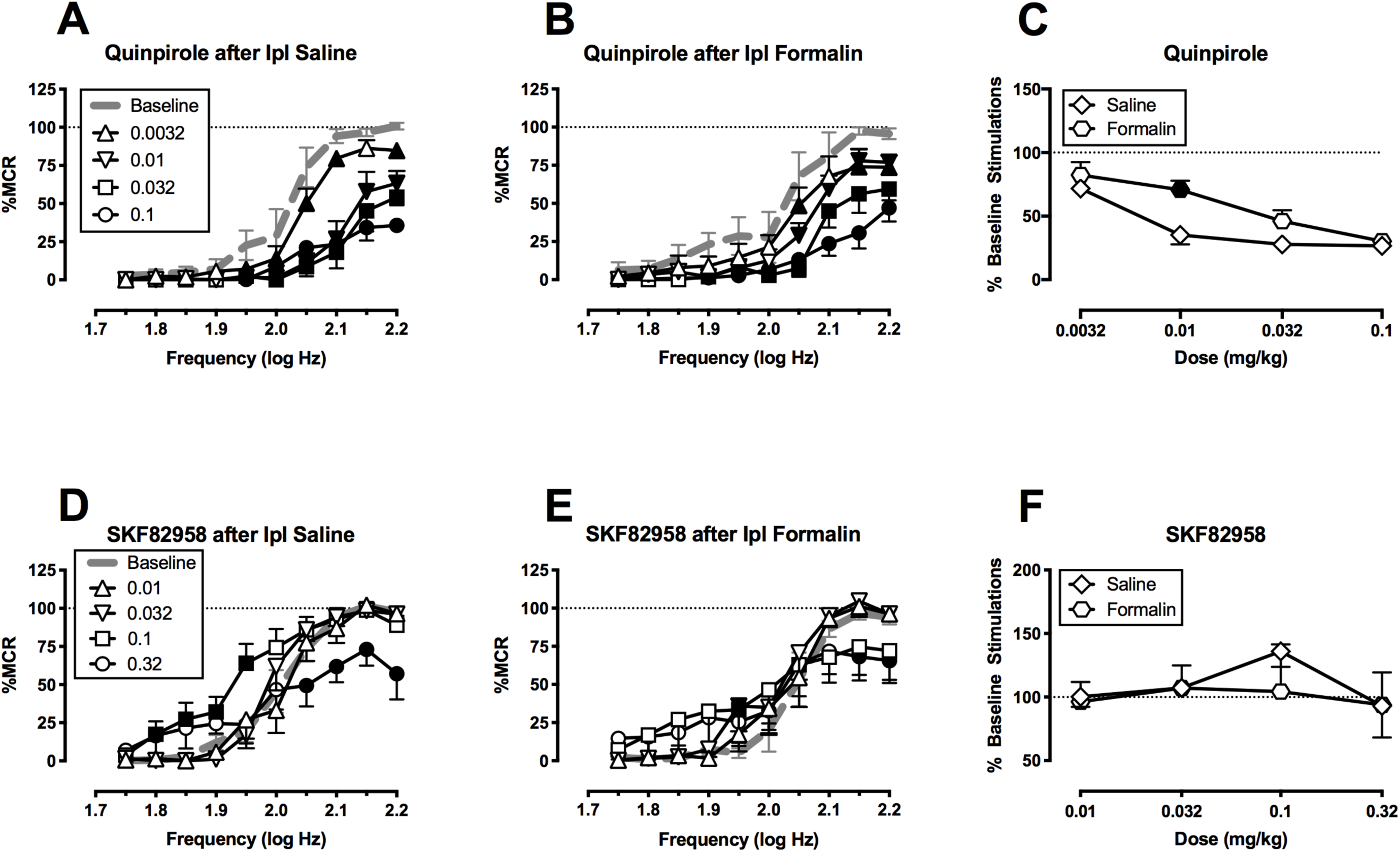
Formalin treatment decreases the potency of D_2_-like but not D_1_-like agonists to modulate ICSS responding. Abscissae: Frequency of electrical brain stimulation in Hz (log scale; A, B, D and E) or dose of drug (C and F). Ordinates: Percent maximum control reinforcement rate (% MCR; A, B, D and E) or percent baseline number of stimulations per component (C and F). Filled symbols (A, B, D and E) show significant differences from baseline as determined by repeated-measures two-way ANOVA followed by the Holm-Sidak post-hoc test, p < 0.05. Filled symbols for panel C show a significant difference between saline and formalin treated rats following a two-way mixed ANOVA and a Holm-Sidak post-hoc test, p < 0.05. All data show mean ± SEM of 6 rats. Statistical results are as follows (only significant interaction results are shown for brevity): (A) significant frequency x dose interaction [F(36, 180) = 10.41, p < 0.0001], (B) significant frequency x dose interaction [F(36, 180) = 3.68, p < 0.0001] (C) significant dose x intraplantar treatment interaction [F(4, 40) = 5.80, p < 0.001], (D) significant frequency x dose interaction [F(36, 180) = 4.78, p < 0.0001], (E) significant frequency x dose interaction [F(36, 180) = 2.32, p < 0.001].

## 4. DISCUSSION

### 4.1 Summary of major findings

The present study examined the effects of bilateral Ipl formalin, a method often used to model of chronic pain in humans, on DA receptor expression and signaling in the mesolimbic DA system and on modulation of ICSS behavior by D_1_-like and D_2_-like receptor agonists. In agreement with previous studies, the overall results of this study indicate that chronic pain manipulations can modulate DA signaling in the mesolimbic system, including changes in D_2_-like receptor expression in NAc. Moreover, three major novel findings were revealed. First, selective reductions in both D_1_R and D_2_R-mediated G-protein signaling were demonstrated within the NAc. These effects were receptor-homologous as suggested by the finding that formalin treatment did not significantly affect activity of the G_s/olf_-coupled adenosine A_2a_ receptor or the G_i/o_-coupled cannabinoid CB_1_ receptor. Second, these effects, along with decreased D_2L_R protein, were seen in NAc core but not the shell. Further, decreased D_2_R expression was specific for the long but not short isoform of the D_2_R, indicating reduced expression of the post-synaptic D_2_R population. Third, the decrease in NAc core D_2_R function and expression was associated with reduced potency of the D_2_-like agonist quinpirole to decrease positively reinforced operant responding in an ICSS procedure; however, behavioral effects of D_1_-like agonist on ICSS were not significantly altered. Altogether, this profile suggests that formalin-induced neuropathy can alter dopaminergic responsivity in the NAc core, with predominant downregulation of D_2_R function.

### 4.2 Formalin treatment attenuated DA receptor signaling and/or expression in NAc core

Several previous preclinical and human functional imaging studies suggest a role for the NAc in both acute and chronic pain (Magnusson and Martin, 2002) (Becerra and Borsook, 2008) (Baliki et al., 2010; Baliki et al., 2013; Becerra et al., 2013; Chang et al., 2014; Schwartz et al., 2014). The NAc can be divided into core and shell regions with evidence showing that each region responds differently to acute thermal pain in humans (Baliki et al., 2013). The present study showed a decrease in quinelorane-stimulated G-protein activation and a specific decrease in the D_2_R receptor long isoform in the NAc core but not shell. In broad agreement with our results, other studies of neuropathic pain in rodents have shown opposing changes in neuron excitability of D_2_R expressing medium spiny neurons in the NAc core and shell (Ren et al., 2016).

In addition to effects on D_2_R expression and activity, the current study also found that AC activity stimulated by the D_1_R-selective agonist SKF82958 was decreased in the NAc core but not shell. To our knowledge, this is the first study to show effects of neuropathy on D_1_R activity in the NAc or any changes in function in the D_1_R-expressing population of medium spiny neurons. D_1_R activation is capable of reversing pain-depressed behavior induced by intraperitoneal acid administration (Lazenka et al., 2017a), and the present findings of reduced D_1_R activity in neuropathic rats suggest a potential for reduced effectiveness of D_1_R agonists. Conversely, A_2a_ receptor activation and D_2_R inhibition were reported to mediate increased pain sensitivity and behavioral depression associated with sleep deprivation (Sardi et al., 2018), suggesting that increased activity of A_2a_/D_2_R-expressing neurons contributes to pain behaviors. Moreover, the decreased ratio of D_1_R to A_2a_ receptor-mediated AC activation coupled with reduced post-synaptic D_2_R expression and activity found in NAc core in the present study suggest that the balance of activity of D_1_R-versus D_2_R-expressing medium spiny neurons would be perturbed by neuropathic pain. Predominant activity of A_2a_-relative to D_1_R-expressing neurons in NAc core could therefore contribute to pain-induced behavioral depression.

The mechanisms underlying desensitization and/or downregulation of D_2_R and D_1_R in NAc core of rats with neuropathy are unclear. A decrease in D_2L_ protein could indicate decreased gene (mRNA) expression, as suggested by prior work on SNI-induced neuropathy (Chang et al., 2014). However, it is also possible that D_2_R desensitization and downregulation occur at the post-translational level, for example in response to increased activation by endogenous DA. In fact, increased DA in the NAc has been reported two weeks after SNI-induced neuropathy in rats (Sagheddu et al., 2015), although decreased DA has also been reported under similar conditions (Wu et al., 2014). The most common mechanism of G-protein-coupled receptor (including DA receptor) regulation in response to agonist occupancy is receptor phosphorylation followed by β-arrestin-mediated desensitization and internalization (Gurevich et al., 2016). In addition, D_2_ receptors interact with G-protein-coupled receptor-associated sorting protein 1 (GASP1), a post-endocytic trafficking protein that promotes trafficking to lysosomes resulting in receptor degradation (Bartlett et al., 2005; Thompson et al., 2010). This mechanism has been shown to mediate D_2_R downregulation in response to repeated cocaine treatment. Therefore, the formalin-induced reduction in D_2L_ receptor protein could have been due to enhanced GASP1-mediated lysosomal degradation of the protein. In contrast, D_1_R protein was unaffected by formalin treatment, consistent with its lack of interaction with GASP1 (Bartlett et al., 2005; Thompson et al., 2010). The reduction in D_1_-like stimulation of AC activity could have been due to receptor phosphorylation and β-arrestin-mediated desensitization. Indeed, D_1_R in striatal neurons interact with β-arrestin2 in an agonist-stimulated manner (Macey et al., 2005), and catechol-containing agonists such as endogenous DA have been shown to effectively recruit β-arrestin2 to the D_1_R (Gray et al., 2018).

There are other potential explanations for reduced D_1_R or D_2_R signaling in NAc core of formalin-treated rats, including increased heteromeric interactions with other G-protein-coupled receptors. For example, D_2_R signaling to G_i/o_ can be reduced by switching to G_s/olf_ or G_q/11_ signaling via heteromomerization with CB_1_ or A_2a_ receptors, respectively (Ferre et al., 2009). However, no significant differences in CB_1_ or A_2a_ receptor activity was seen between formalin- and saline-treated rats in the present study. Nonetheless, mechanisms including homologous or heterologous receptor regulation and heteromerization are not generally thought to be mutually exclusive, so future studies will be required to tease out the contribution of these molecular processes in DA receptor adaption in neuropathic pain states.

### 4.3 Formalin treatment decreased D_2_-like agonist potency in intracranial self-stimulation

Ipl formalin is well-established in preclinical studies as a chronic-pain stimulus (Fu et al., 2001; Grace et al., 2014), and we reported previously that Ipl formalin served as a chemical neuropathic stimulus that produced a small but significant decrease in ICSS that was sustained for up to 14 days(Leitl and Negus, 2016; Leitl et al., 2014b). This sustained ICSS depression served as the rationale in the present study to examine dopamine receptor function and expression 14 days after Ipl formalin, however the present study failed to replicate this Ipl formalin-induced ICSS depression. Other treatments that produce neuropathy in rats, such as spinal nerve ligation or chemotherapy administration, are also generally ineffective to decrease ICSS and other forms of positively reinforced operant behavior in rats (Ewan and Martin, 2011; Legakis et al., 2018; Okun et al., 2016). Taken together, these results suggest that Ipl formalin and other neuropathy manipulations are at best weakly and inconsistently effective to depress ICSS in rats, and results of the present study regarding dopamine receptor function and expression may not be related to pain-related behavioral depression. Factors that underlie the resistance of operant behavior to neuropathic pain-related depression remain to be determined, but it is possible that compensatory increases in mesolimbic dopaminergic activity may have occurred by two-weeks post injury as previously reported using the rat SNI model of neuropathic pain (Sagheddu et al., 2015). Such an adaptation could potentially underlie both the lack of ICSS depression and the DA receptor desensitization and downregulation observed in NAc core. Future studies could examine individual differences among subjects in response to neuropathy-inducing injury by comparing ICSS responses and function/expression of DA receptors, for example by comparing different strains or using recombinant inbred panels of animals.

Despite the lack of effect on baseline ICSS responding, it remained possible to examine effects of formalin treatment on sensitivity to the effects of D_1_-like and D_2_-like receptor agonists. We reported previously that quinpirole and other D_2_-like receptor agonists produce dose-dependent decreases in ICSS, whereas D_1_-like agonists produce a mixed profile with stimulation by low doses and primarily depression by high doses (Lazenka et al., 2016). In the present study, formalin treatment decreased the potency of the D_2_-like receptor agonist quinpirole to decrease ICSS, which agrees with the decrease in D_2_R-stimulated G-protein activation seen in the NAc core. However, it is not clear if this effect is mediated by the D_2_R long or short isoforms or dopamine D_3_ receptors since both quinpirole and quinelorane can activate all of these receptors. The effectiveness of D_2_-like receptor agonists to depress ICSS is likely due to decreased DA release through activation of the D_2S_ and dopamine D_3_ autoreceptors (Gilbert et al., 1995; Khan et al., 1998; Lindgren et al., 2003; Sokoloff et al., 1990; Usiello et al., 2000). Decreased expression and signaling of the D_2L_R may also play a role in decreased ICSS effects of the D_2_-like agonist, although the potential mechanism is less clear. It is also possible that some of the decrease in D_2_-like agonist-stimulated G-protein activation in NAc core of formalin-treated rats was actually due to decreased signaling of D_2s_ or D_3_ receptors. In agreement with this interpretation, agonist-stimulated G-protein activation was reduced by 38% in formalin-relative to saline-treated rats, whereas D_2L_R protein was only reduced by 18%. While this difference in magnitude of effect could be the result of desensitization of a portion of the remaining D_2L_R, it could also be due to a greater population of D_2_-family receptors (including D_2S_ and D_3_) being desensitized.

Despite a decrease in D_1_R-stimulated AC activity in the NAc core of formalin-treated rats, there was no significant change in D_1_R expression Moreover, there was no change in the effects of a D_1_-like agonist on ICSS responding, although the intermediate dose of 0.1 mg/kg SKF82958 generally produced weaker ICSS facilitation in formalin-than saline-treated rats. One explanation for this lack of effect may be the biphasic nature of D_1_-like agonist effects on ICSS, making subtle changes difficult to detect. Furthermore, it may be that the reinforcing effects of ICSS are more reliant on D_1_R expression and function in the NAc shell than core (Cheer et al., 2007), which would be consistent with our observation of no differences in D_1_R-stimulated AC activity in the shell. The overall results of these studies indicate that decreased D_2_-like receptor signaling in the NAc core of formalin-treated rats corresponded with decreased potency to depress ICSS by a D_2_-like agonist, whereas decreased D_1_-like receptor signaling in NAc core did not correspond with D_1_-like agonist effects on ICSS. However, the underlying mechanisms of these behavioral effects are likely to be complex.

### 4.4 Conclusions

In conclusion, this study evaluated DA receptor expression and function in a subset of mesolimbic dopaminergic brain regions associated with motivated behavior in a chemically-induced neuropathy model in rats. The findings indicate attenuation of both D_1_-like and D_2_-like receptor signaling in the NAc core but not in other regions examined, coupled with a selective reduction in D_2L_R expression in NAc core. These findings were associated with decreased potency of D_2_-like but not D_1_-like agonists to modulate ICSS responding. Moreover, the decreased ratio of D_1_ to A_2a_ receptor-mediated activation of AC, coupled with downregulated D_2_R function, could shift the balance of activity to increase that of D_2_/A_2a_-expressing relative to D_1_-expressiong neurons in NAc core. These findings further support the concept that chronic neuropathy can produce lasting perturbations in mesolimbic dopaminergic system function and point to NAc core as a key region in these allostatic adaptations.

## 5. ACKNOWLEDEMENTS

Supported by National Institutes of Health grant R01-NS070715

