## Supplemental Material for "Attenuated Dopamine Receptor Signaling in Nucleus Accumbens Core in a Rat Model of Chemically-Induced Neuropathy"

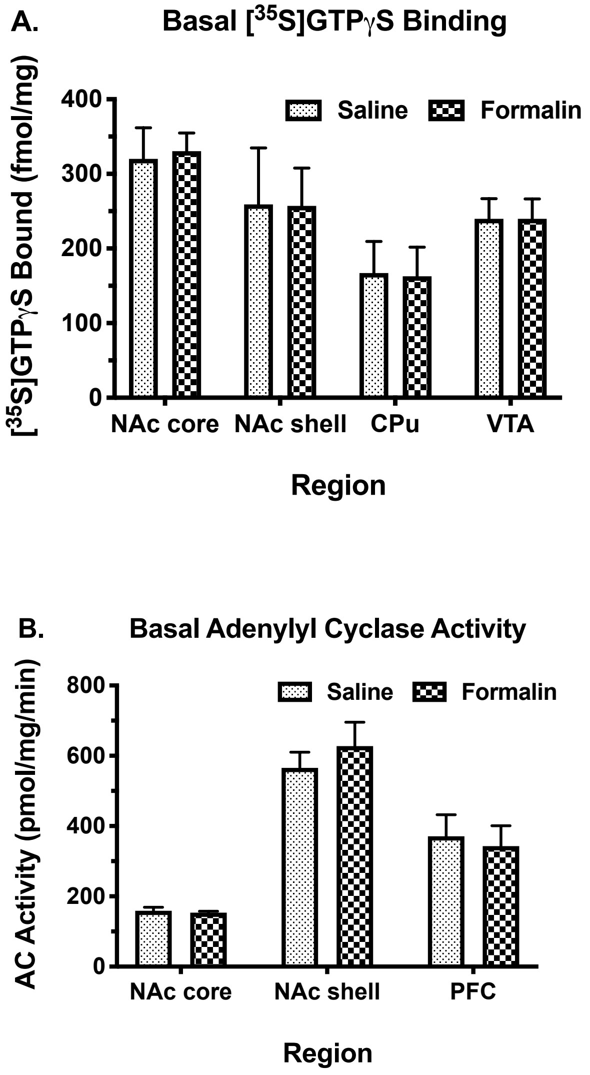


**Supplemental Figure 1.** Basal [$^{35}$S]GTPγS binding and adenylyl cyclase (AC) activity in brain regions of rats treated with intraplantar formalin or saline. Data are mean ± SEM (n = 5-6) of basal [$^{35}$S]GTPγS binding (A) and AC activity (B). Two-way ANOVA of basal [$^{35}$S]GTPγS binding showed a main effect of region [$F(3, 40) = 5.007$, $p = 0.005$] but not of formalin treatment [$F(1,40) = 0.001$, $p = 0.975$], nor was there an interaction [F(3,40 = 0.011, $p = 0.998$]. Two-way ANOVA of basal AC activity showed a main effect of region [$F(2, 27) = 38.93$, $p < 0.0001$] but not of formalin treatment [$F(1,27) = 0.055$, $p = 0.816$], nor was there an interaction [$F(2,27) = 0.442$, $p = 0.648$].

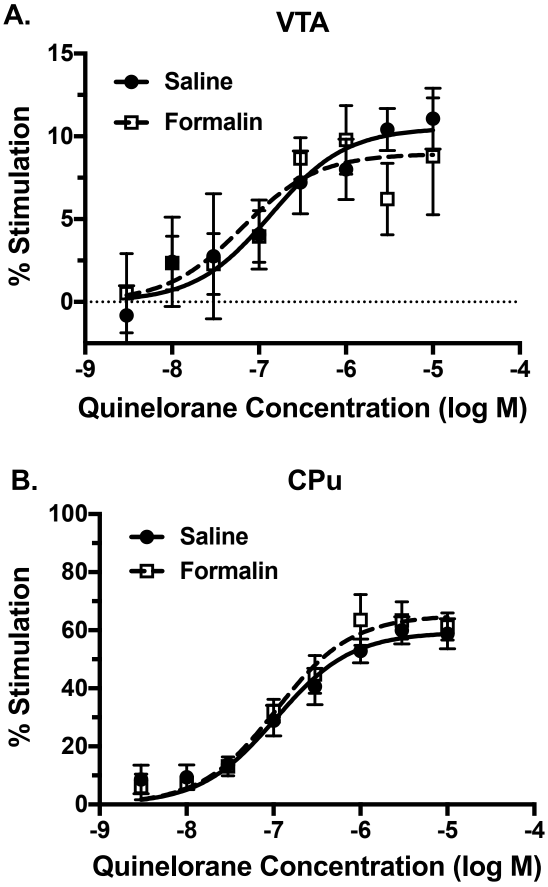


**Supplemental Figure 2.** $D_2$-like receptor-stimulated G-protein activity in ventral tegmental area (VTA) and caudate-putamen (CPu) of rats treated with intraplantar formalin or saline. Concentration-effect curves were conducted for stimulation of [$^{35}$S]GTP$\gamma$S binding by quinelorane in membranes from VTA (A) or CPu (B). Data are mean ± SEM of % stimulation of [$^{35}$S]GTP$\gamma$S binding (n = 5-6). In VTA, two-way ANOVA showed a significant main effect of quinelorane concentration [F(7,63) = 6.050, p < 0.0001], but there was no effect of formalin treatment [F1,63) = 0.139, p = 0.710) nor an interaction [F(7,63) = 0.546, p = 0.796]. In CPu, two-way ANOVA showed a significant main effect of quinelorane concentration [F(7,29) = 60.48, p < 0.0001] but no effect of formalin treatment [F(1,29) = 2.01, p = 0.167] nor an interaction [F(7,29) = 0.593, p = 0.756].

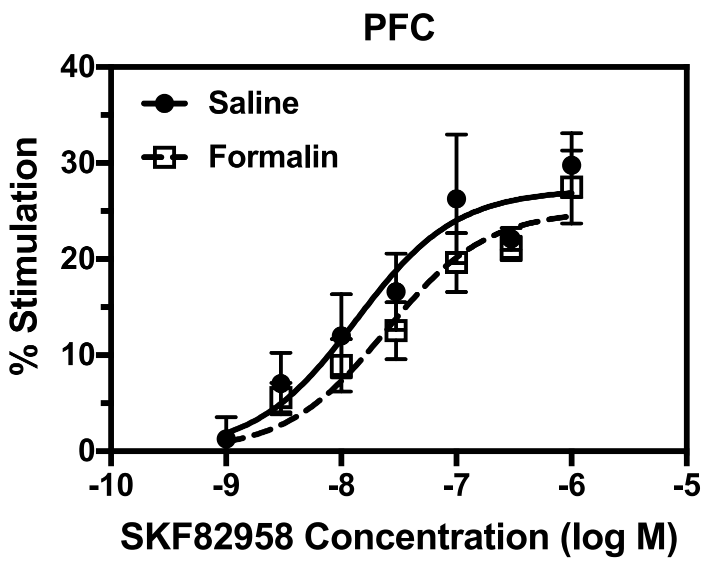


**Supplemental Figure 3.** $D_1$-like receptor-stimulated AC activity in PFC of rats treated with intraplantar formalin or saline. Concentration-effect curves were conducted for stimulation of AC activity by SKF82958. Data are mean ± SEM of % stimulation of AC activity (n = 4-5). Two-way ANOVA showed a main effect of SKF82958 concentration [$F(6,47) = 20.09$, $p < 0.0001$] but not of formalin treatment [$F(1,47) = 3.063$, $p = 0.087$] nor an interaction [$F(6,60) = 0.176$, $p = 0.920$].

A.

B.

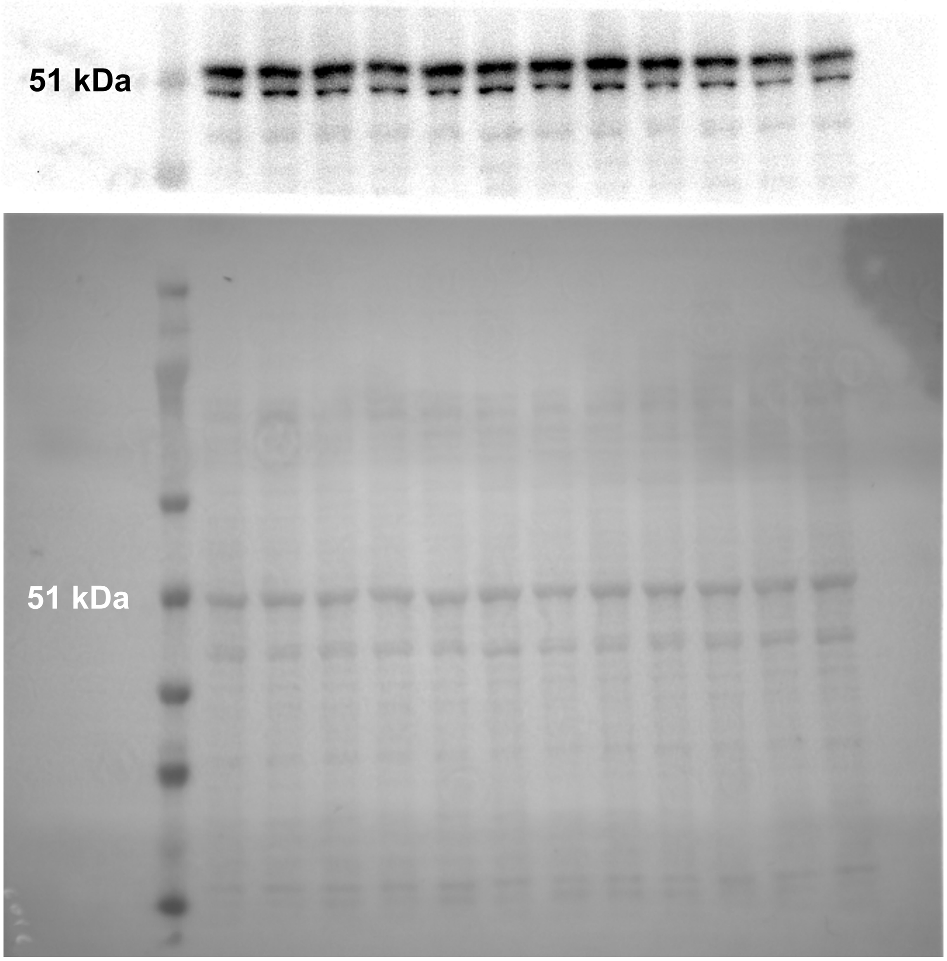


**Supplemental Figure 4.** $D_2$ receptor immunoreactivity in NAc core of rats treated with intraplantar formalin or saline. Images are $D_2$ receptor immunoblot (A.) and corresponding Ponceau stain (B). Lanes are: 1, M.W. marker; 2, 4, 6, 8, 10, 12, saline; and 3, 5, 7, 9, 11, 13, formalin. Graphs of densitometry results are shown Figure 3, panels C and E.

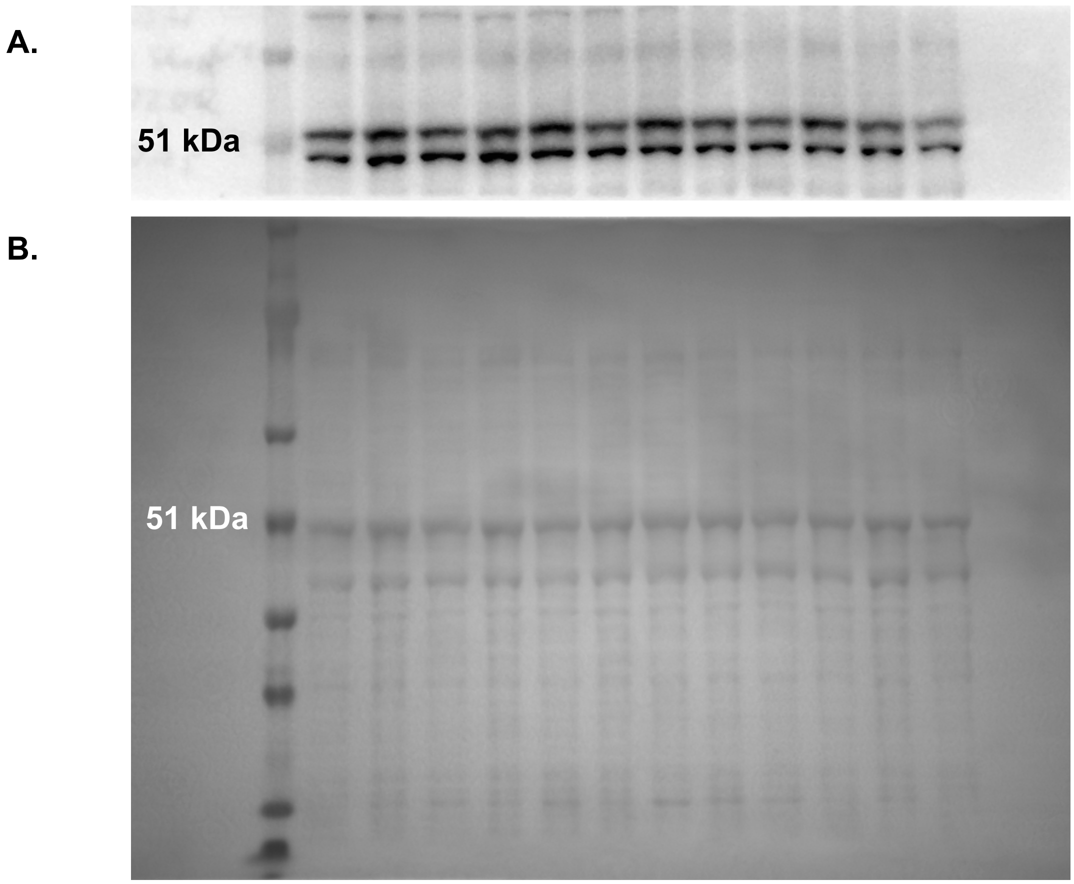


**Supplemental Figure 5.** $D_2$ receptor immunoreactivity in NAc shell of rats treated with intraplantar formalin or saline. Images are $D_2$ receptor immunoblot (A.) and corresponding Ponceau stain (B). Lanes are: 1, M.W. marker; 2, 4, 6, 8, 10, 12, saline; and 3, 5, 7, 9, 11, 13, formalin. Graphs of densitometry results are shown Figure 3, panels D and F.

A.

B.

C.

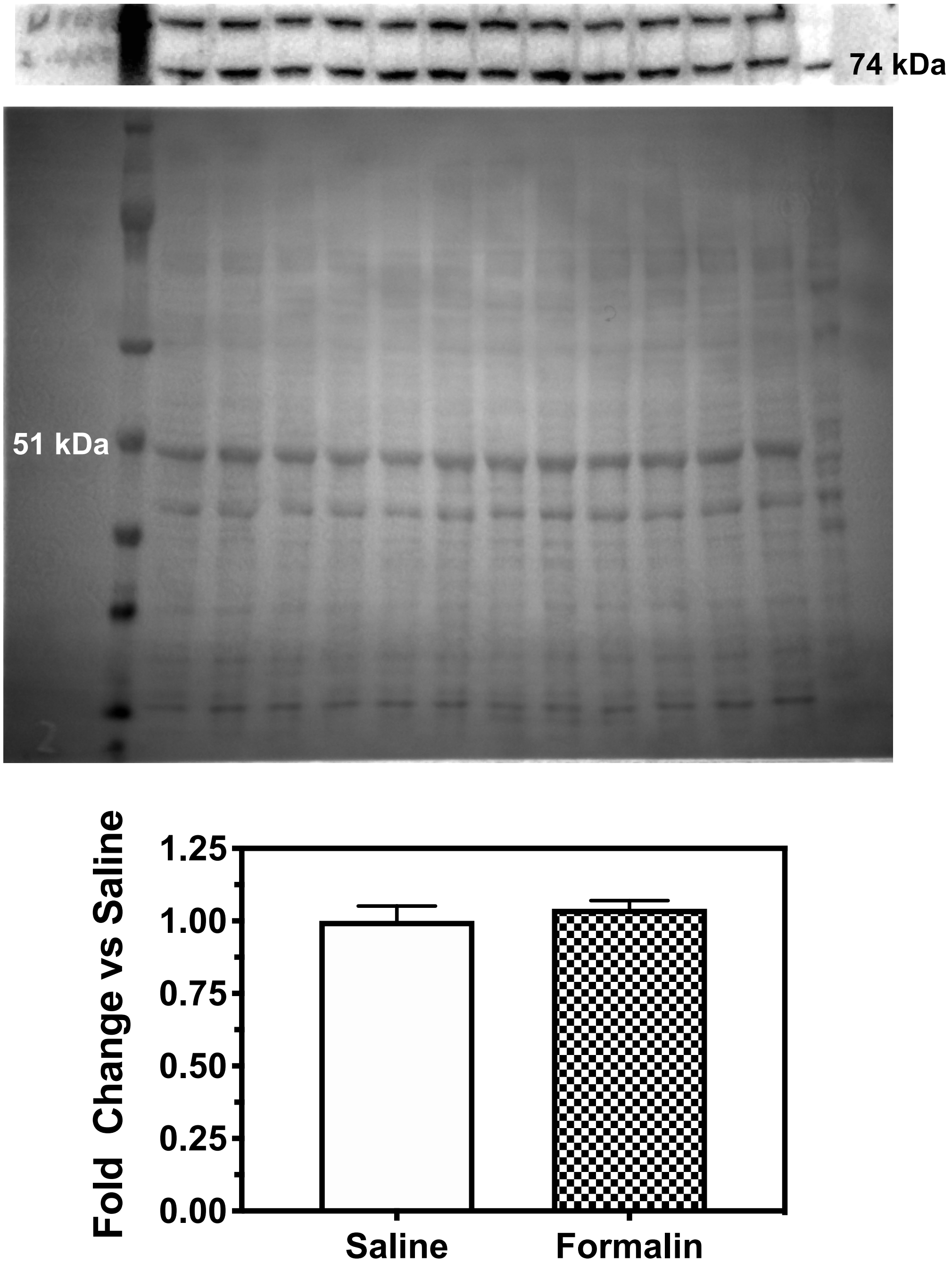


**Supplemental Figure 6.** $D_1$ receptor immunoreactivity in NAc core of rats treated with intraplantar formalin or saline. Images are $D_1$ receptor immunoblot (A.), corresponding Ponceau stain (B) and densitometry results (C) (n = 6) (t = 0.719, p = 0.489; n = 6). Lanes are: 1, 14, M.W. marker; 2, 4, 6, 8, 10, 12, saline; and 3, 5, 7, 9, 11, 13, formalin.

A.

B.

C.

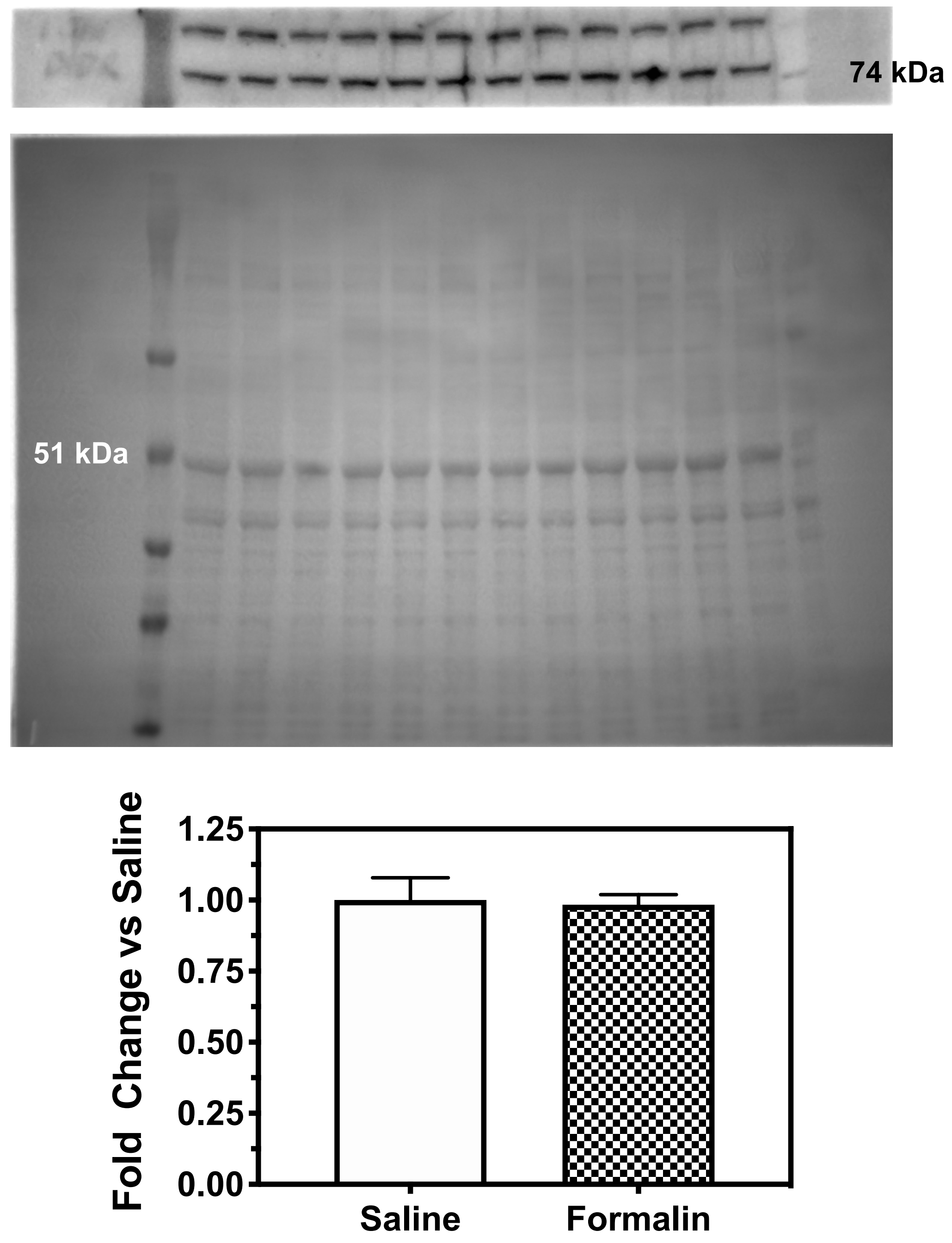


**Supplemental Figure 7.** $D_1$ receptor immunoreactivity in NAc shell of rats treated with intraplantar formalin or saline. Images are $D_1$ receptor immunoblot (A.), corresponding Ponceau stain (B) and densitometry results (C) ($t = 0.0.164$, $p = 0.874$; $n = 4\text{-}6$). Lanes are: 1, 14, M.W. marker; 2, 4, 6, 8, 10, 12, saline; and 3, 5, 7, 9, 11, 13, formalin.
